# TGF-β promotes microtube formation in glioblastoma through Thrombospondin 1

**DOI:** 10.1101/2021.02.22.431443

**Authors:** Justin V. Joseph, Capucine R. Magaut, Simon Storevik, Luiz H. Geraldo, Thomas Mathivet, Md Abdul Latif, Justine Rudewicz, Joris Guyon, Matteo Gambaretti, Frida Haukas, Amalie Trones, Lars A. Rømo Ystaas, Jubayer A. Hossain, S Sandra Ninzima, Wenjing Zhou, Tushar Tomar, Barbara Klink, Lalit Rane, Bronwyn K. Irving, Joanne Marrison, Peter O’Toole, Heiko Wurdak, Jian Wang, Zhang Di, Frank Winkler, Frank A. E. Kruyt, Andreas Bikfalvi, Rolf Bjerkvig, Thomas Daubon, Hrvoje Miletic

## Abstract

Microtubes (MTs), cytoplasmic extensions of glioma cells, are important cell communication structures promoting invasion and treatment resistance through network formation. MTs are abundant in chemoresistant gliomas, in particular glioblastomas (GBMs), while they are uncommon in chemosensitive IDH-mutant and 1p/19q co-deleted oligodendrogliomas. To identify potential signaling pathways involved in MT formation we performed a bioinformatics analysis of TCGA data showing that the TGF-β pathway is highly activated in GBMs compared to oligodendroglial tumors. In particular we observed that signaling pathways involved in extracellular matrix organization are differentially expressed between these tumor entities. Using patient-derived GBM stem cell lines, we demonstrated that TGF-β1 stimulation promotes enhanced MT formation and communication via Calcium signaling. Inhibition of the TGF-β pathway significantly reduced MT formation and its associated invasion *in vitro* and *in vivo*. Downstream of TGF-β, we identified thrombospondin 1 (TSP1) as a potential mediator of MT formation in GBM through SMAD activation. TSP1 was upregulated upon TGF-β stimulation and enhanced MT formation, which was inhibited by TSP1 shRNAs *in vitro* and *in vivo*. In conclusion, TGF-β and its downstream mediator TSP1 are important mediators of the MT network in GBM and blocking this pathway could potentially help to break the complex MT driven invasion/ resistance network.

## Introduction

Malignant gliomas, the largest group of primary intracerebral tumours, are one of the most difficult-to-cure cancers. The outcome has not improved significantly during the last decades, despite considerable advances in our understanding of the molecular pathogenesis and the improvement of surgical techniques as well as radio- and chemotherapy. For glioblastoma (GBM), the most malignant form of glioma, the median survival time is approximately 15 months [28]. Thus, a basic understanding of the disease is necessary to develop novel and effective therapies, which are urgently needed.

In 2015, Osswald and colleagues discovered the existence of microtubes (MTs), which are cytoplasmatic extensions of GBM cells [26]. In this hallmark paper, the authors showed that MTs form a connective network leading to increased invasion and promoting resistance to radiotherapy. Growth Associated Protein 43 (GAP43), a protein involved in neurite sprouting, was identified as an important mediator involved in the structural development of MTs. Intercellular calcium waves were detected as a way of communication through the MT network and Connexin 43, a major gap junction protein, was found to be functionally involved [26]. The MT network was observed extensively in GBM and also in astrocytic tumors in general, however, was scarce in oligodendroglioma. These two tumor types can be distinguished at the molecular level by IDH mutation and 1p/19q co-deletion, which are only present in oligodendroglioma [22]. In contrast to GBMs and astrocytomas, oligodendroglioma patients show remarkable responses to radio- and chemotherapy and therefore have a much better prognosis [31]. Thus, to identify mechanisms of MT formation in GBM and to unravel how MTs promote cell-to-cell communication might ultimately lead to a better understanding of therapeutic resistance and new ways of GBM treatment. Interestingly, a recent study showed that network formation of MTs was enhanced in the resection cavity after neurosurgery in experimental models [34]. MT formation in this context might partly explain why recurrent tumors often develop close to the resection site. Recently, it was shown that glioma cells and neurons are connected through so-called neurogliomal synapses that are located on MTs indicating another layer of resistance through connection with normal cells [32]. Molecular mediators driving the MT network identified so far such as GAP43 and tweety-homolog 1 (TTYH1) [17] are also important in normal physiology of the CNS. Thus, the identification of more tumor specific pathways that drive MT formation is of great interest in order to develop targeted therapies.

We recently showed by RNA sequencing of patient-derived xenograft (PDX) tissue from laser capture micro-dissected invasive and central tumour areas that the matricellular protein thrombospondin-1 (TSP1 or THBS1) was one of the most up-regulated genes in infiltrative areas of GBM. TSP1 is a large trimeric calcium-binding molecule, which binds to diverse ligands and receptors [1]. The TGF-β1 canonical pathway transcriptionally regulates TSP1 expression, through SMAD3 binding to the *THBS1* promoter. TSP1 silencing inhibited cell invasion *in vitro* and *in vivo*, also in combination with anti-angiogenic therapy [10]. Interestingly, TSP1 was also found to be regulated by calcium flux in several cancers [13]. In the present study we showed that TGF-β1 stimulation of GBM stem-like cell cultures promotes MT formation and that TSP1, a protein highly upregulated in GBM, is an important mediator of MT formation downstream of TGF-β.

## Materials and Methods

### Ethics statement

Patient material was obtained from surgeries performed at the Haukeland University Hospital (Bergen, Norway). Written consent was obtained from patients with procedures that were approved for the projects (project numbers 013.09 and 151825) by the Regional Ethics Committee (Bergen, Norway). In vivo experiments were conducted in accordance with the European Community for experimental animal use guidelines (L358-86/609EEC) with protocols approved by the Ethical Committee of INSERM (n°MESRI23570).

### TCGA data and GMT files

For the ‘The Cancer Genome Atlas’ (TCGA) dataset, RNAseqV2 median normalized data (RNA Seq V2 RSEM) of the TCGA GBM Multiforme cohort and associated clinical data were downloaded from cBioPortal (https://www.cbioportal.org/study/summary?id=gbm_tcga). RNAseqV2 median normalized data (RNA Seq V2 RSEM) of the TCGA Lower Grade Glioma cohort (LGG) and associated clinical data were downloaded from cBioPortal (https://www.cbioportal.org/study/summary?id=lgg_tcga). The associated IDH_1P19Q_SUBTYPE and IDH_CODEL_SUBTYPE status for samples were downloaded from cBioPortal (https://www.cbioportal.org/study/summary?id=lgggbm_tcga_pub). 513 LGG and 143 GBM primary tumor samples were used resulting in 656 fully annotated samples. Of note, only 654 of the 656 have Overall Survival (OS) information. Gmt files (v6.2) were download from GSEA (https://www.gsea-msigdb.org/gsea/downloads.jsp). Differential expression and enrichment analysis were done with R language in R studio (R v3.5.2 [30], R studio v1.1.463 [29]).

### DEG analysis and visualisation

We used DESeq2 (v.1.22.2)[23] R package to perform differential expression analysis. Preprocessing and negative Binomial Generalized Linear Models fitting were applied to pre-transformed expression values of TCGA RNAseq data set. Wald statistics was then performed for 2 group differences (IDH_1P19Q_SUBTYPE; non-codel vs codel) and likelihood ratio test (LRT) for 3 group differences (IDH_CODEL_SUBTYPE; IDH wt, IDH mut codel and IDH mut codel).

To select the most robust differentially expressed genes (DEGs), we considered a gene as significantly differentially expressed if its absolute log2FC was greater than or equal to 1 and its -log10 (padjust) was in the last quartile of the -log10 (padjust) distribution of all gene expression for the 2 differential analyses. For the visualization of our results (boxplot and heatmap), we used variance stabilizing transformations (vst) that remove the dependence of the variance on the mean (function vst of DESeq2 with the argument blind set to TRUE). Vst values were centered and scaled for the pheatmaps of the TGF-β responsive genes.

### Enrichment analysis

Enrichment analysis was performed with the clusterProfiler (v.3.10.1 [37], enricher function) R package using the c2.cp.reactome, c2.cp.kegg and c5.all (Gene Ontology) gene sets from the molecular signatures database [20]. We used biomaRt (v.2.38.0)[11] R package to translate Entrez Gene ID to gene Name. For a given pathway, its log2FC was computed as the mean expression of all the DEGs it contains. Its zscore was then computed as previously described [33].

To build the heatmap focusing on TGFB1 or TGFB2, we selected the top 50 DEGs sharing the most GO terms with TGFB1 or TGFB2 and the associated top 20 GO terms containing the most of these 50 genes. Heatmap were plotted with pheatmap (v.1.0.1) R package.

### Cell culture

The patient derived GSC lines P3[12], GG6 and GG16[16] were cultured in Neurobasal Medium, B27 supplement, Glutamax (NBM; Life Technologies), FGF2 (20ng/ml), and EGF (20ng/ml; Peprotech) in non-treated culture flasks (Nunc). U87, U251 and 293T cells were obtained from the American Type Culture Collection (VA, USA) and maintained in Dulbecco’s modified eagle medium (DMEM) supplemented with 10% fetal calf serum (FCS) and 1% glutamine. Medium was changed twice a week. All cell lines were grown at 37°C in a humidified atmosphere of 5% CO_2_.

### Matrigel coating

Growth factor reduced Matrigel (Corning) was thawed over ice and resuspended in appropriate volume of neurobasal medium or DMEM medium without any added supplements. Matrigel was diluted to a concentration of 0.2mg/ml and 1 ml or 0.2 ml of this diluted solution was added to each well of a 6-well or a 4-well plate, respectively, and incubated at 37°C for 30 minutes. After this incubation step, matrigel was aspirated and the cells were seeded immediately and incubated at 37°C in a humidified atmosphere of 5% CO_2_.

### Lentiviral shRNA constructs and production

shRNA target sequences for TSP1 [10] and GAP43[26] have been described previously. The inducible shRNA construct for TGFBRII Lentiviral vectors was designed by inserting a target sequence (GCTTCTCCAAAGTGCATTATG) into doxycycline-dependent pLKO1 vectors, then lentivirus were produced in HEK293T cells using CaCl_2_ transfection and packaging plasmids as described previously. GBM cells were then transduced as described previously [10].

### Western blots

Cultured cells were harvested, washed with cold PBS and lysed with appropriate volume of M-PER^™^ mammalian protein extraction agent (Thermo Fisher Scientific) supplemented with 1% protease inhibitor (Thermo Fisher Scientific) and 1% phosphatase inhibitor (Thermo Fisher Scientific) for 30 min on ice. Next, the suspension was centrifuged for 10 min at 14000 r.p.m. at 4°C and the supernatant was taken for determining protein concentrations using a Direct Detect^®^ Infrared Spectrometer (Merck). Whole lysate was used for subsequent analysis. Twenty micrograms of protein were loaded in each well. Protein was run on SDS/PAGE by using NuPage precast gels (Life Technologies, Carlsbad, CA). After blotting, the nitrocellulose membrane was blocked for 30 min at room temperature and incubated overnight at 4°C in buffer (TBS with 0.1% Tween 20, 5% milk powder) containing the primary antibody. Primary antibodies used were anti-Nestin [1:2000, Sigma-Aldrich (MAB5326)], Anti-GAP43 [1:10000, Abcam (ab75810)], Anti-F-actin [1:500, Abcam (ab130935)], Anti-Phospho-SMAD2 [1:1000, Cell Signaling (3108)], Anti-TGFβ RII [1:250, Santa Cruz Biotechnology (sc-17791)], anti-TSP1 [1:500, Invitrogen], anti α -Tubulin [1:5000, Sigma], anti-Vinculin [1:2000, Invitrogen], Anti-Phospho-MAPK [1:2000, Cell Signaling (4370)], Anti-Phospho-AKT [1:1000, Cell Signaling (9271)] anti-GAPDH [1:10000, Invitrogen (PAI-16777]. The primary antibody was detected by using an HRP-conjugated goat anti-rabbit/mouse secondary antibody (Immunotech, Fullerton, CA) diluted 1:2500. Membranes were probed with GAPDH, tubulin, or vinculin antibody to confirm equal loading.

### Immunofluorescence staining

Cells cultured on matrigel (Corning)-coated cover slips were fixed for 10 min using 4% formaldehyde or 100% methanol. After 3 times washing with cold PBS, cells were permeabilized with 0.1% Triton (Sigma-Aldrich) in PBS, washed again with PBS followed by a blocking step for 1 hr with PBS + 0.1% Tween-20 (Sigma-Aldrich), 2% BSA (PAA Laboratories GmbH, Germany) and 1:50 dilution of normal goat serum (Dako Denmark A/S, Denmark). Subsequently, cells were incubated with the indicated primary antibodies at room temperature for 1.5 hrs. Primary antibodies used: anti-Nestin [1:250, Sigma-Aldrich (MAB5326)], anti-GAP43 [1:500, Abcam (ab75810)], anti-F-actin [1:250, Abcam, (ab130935), anti-Phospho-SMAD2 [1:400, Cell Signaling, (18338)]. After 3-times washing with PBS, slides were incubated for 1 hr with the appropriate secondary antibodies: Goat Anti-Mouse Alexa647 [1:400, Abcam, (ab150115)], Goat Anti-Rabbit Alexa488 [1:400, Abcam, (ab150077)]. Thereafter, the coverslips were washed three times in PBS and mounted using ProLong^™^ Gold Antifade Mountant with DAPI. Cells were examined by fluorescence microscopy (Leica DM6000, Leica Microsystems GmbH, Mannheim, Germany) and images were captured using Leica DFC360 FX camera.

### Immunohistochemistry

Paraffin embedded formalin-fixed tissue sections were placed in xylene bath for 2×3 minutes, absolute ethanol 2×3 minutes, 96% ethanol 2×2 minutes and finally in distilled water for 30 seconds for removal of paraffin and rehydration. Epitope retrieval was performed by heating the sections at 99°C for 20 minutes in 10 mM citrate buffer at pH 6.0. The sections were incubated with primary antibodies in TBS/1%BSA over night at 4°C. The following primary antibodies were used: monoclonal human-specific anti-nestin antibody [1:250, Sigma-Aldrich (MAB5326)] and anti-Phospho-SMAD3 [1:100, Abacam (ab52903)]. A biotinylated goat-anti-mouse antibody (Vector Laboratories) was used as secondary antibody (dilution 1:200) for 1 h at room temperature followed by ABC-complex incubation for 30 min. Sections were developed with 3′3-diaminobenzidine (DAKO Cytomation), following the manufacturer’s instructions.

### Scanning Electron microscopy

Scanning electron microscopy (SEM) was used to investigate the MTs in greater detail. GBM cells were seeded on Matrigel coated 12mm coverslips placed in a 12-well plate. 24 hrs post seeding the cells, TGF-β1 was added at a concentration of 10ng/ml. The samples were then first fixed with 2.5 % glutaraldehyde in Sørensens buffer (pH 7.4) for 2 h and rinsed before they were treated with 1 % osmium (OsO4) for 1 h. After serial stepwise ethanol dehydration and critical point drying, samples were mounted on aluminium stubs and sputter coated with gold palladium. MTs were observed under a Field Emission SEM (Jeol JSM-7400 F, Jeol, Tokyo, Japan) and micrographs were recorded at an accelerating voltage of 5.0 kV.

### MT quantifications

MT length was quantified using the NIH Image J software. DAPI and nestin co-stained 20x magnified microphotographs from three independent experiments were imported and ™ length was measured manually using freehand measurement tool after setting scale at 20x magnification (438 pixels = 200 um). For each experimental condition, a minimum of 80-120 tubes showing a direct connection between two tumor cells, were included for measurement. All measured data were exported into excel and GraphPad PRISM 8.1.2 for further calculation of statistical significance.

For the MT quantifications on patient and xenograft sections, whole-slide Images (WSI) were acquired using a Hamamatsu C12000-02 Slide Scanner (Hamamatsu Photonics, Hamamatsu City, Japan), and analyzed using Fiji software. Automated quantifications of MTs was done by first using color deconvolution on Nestin-stained sections to isolate the DAB signal, then enhancing tubular structures using the Jerman enhancement filter ^[15]^. Blob-like structures were enhanced using the same filter with different parameters, and in addition to the perimeter around nuclei, these were removed. The resulting image was then binarized by thresholding with a fixed threshold and skeletonized to make a simplified network. Short branches were removed from the resulting network using the Analyze Skeleton plugin for Fiji ^[3]^. Finally, the total length of the network was quantified using the same plugin.

pSMAD3 quantification was similarly done by first using color deconvolution to separate DAB and haematoxylin channels, before smoothing the output images with, respectively, median and Gaussian filters. The images were then binarized by thresholding, using a fixed threshold for pSMAD3 to define positive cells, and an adaptive threshold for hematoxylin. This was followed by separation into individual cells using watershed and connected components segmentation, which were further filtered on size, with the minimum area being 12.5 µm^2^. The total number of cells, including the number of pSMAD3+ cells, was finally quantified. To correlate findings from quantification of MTs and pSMAD3 in patient tissue, first all medium-density and invasive areas were marked in sections stained for nestin. This partly because of the hypothesized importance of MTs in the invasive process, and partly because picking out tubular structures in the high-density environment of the tumor core is very challenging in a 2D image. From these chosen areas, a sample of 20 smaller square ROIs was made using an evenly spaced grid. To ensure that the areas quantified for pSMAD3 spatially corresponded to the areas quantified for MTs, we used Fiji’s Register Virtual Slices plugin. Taking the nestin-stained sections as the reference, we performed registration of the nestin and pSMAD3 images in low magnification using an Affine model. This model was finally used to generate the 20 quantification ROIs for pSMAD3 quantification from the original MT ROIs.

### RNA sequencing and analysis

A total of 3 samples were assigned in each of the control (untreated) and treatment (TGF-β_48 hours) groups. Samples were processed for sequencing at Macrogen Inc, Korea where library preparation, sequencing and quality control were performed. Briefly, TruSeq Stranded mRNA sample preparation kit (Illumina) was used for the preparation of the library for RNA-seq data analysis. Poly-A containing mRNA molecules were purified from the total RNAs using poly-T oligo attached magnetic beads. Following purification, RNA fragments were used as templates for first-strand cDNA synthesis by reverse transcription with random hexamers. Upon second-strand cDNA synthesis, double-stranded cDNAs were end-repaired and adenylated at the 3′ ends followed by subsequent ligation of the adapter. The sequencing library was generated by PCR and used to produce the clusters thereafter sequenced on NovaSeq 6000 System (Illumina). Each sample was sequenced in a separate flow cell lane, producing 29-45 M paired-end reads, with a final length of 101 bases.

Illumina’s pipeline was used to generate the raw FASTQ files. *Bowtie2* was used to align the FASTQ files to the reference human genome, hg38, using default parameters. Aligned BAM files were indexed and co-ordinate sorted by Picard SortSam for downstream convenience. Read counts for gene expression were obtained using *FeatureCounts* function [19]. Resulting read count data were then analysed by pairwise comparisons of the groups in R statistical environment (https://cran.r-project.org) using DESeq package [2]. Resulted features with the padj values of less than[0.05 were considered significant. Volcano plot was used to visualize the differentially expressed genes (DEGs) between control and treatment groups. Positive log2 fold change values were used to sort up regulated genes followed by the visual representation using Venn diagram (R package, version 1.6.20). ClusterProfiler [37] was used for the functional enrichment analysis as “biological process over representation” of the up regulated genes.

### GBM-brain organoid co-culture *ex vivo* invasion assay

For this assay, the preparation and culture of 18-day fetal brain organoids have been described in our previous work [6]. After 21 days in culture, the differentiation of cells in the brain organoids was completed and the organoids were ready to be confronted with tumour cells. GFP-GG6 or GFP-GG16 cells were seeded into 96-well plates for 4 days to generate tumour spheroids and then co-cultured with mature brain organoids for 72 h. Confocal microscopy was used to capture images of tumour cell invasion (Leica TCS SP8). ImageJ software (https://imagej-nih-gov.proxy.insermbiblio.inist.fr/ij/, USA) was used for invasion analysis.

### In vitro Calcium imaging

Calcium signal spread across cells was carried out as previously described [9]. Fluo-3-AM (Thermo Fisher Scientific, F23915) was added to the cells and incubated for 30 min in the dark at room temperature. The cells were then washed with PBS and kept in the dark for an additional 30 min. Live-cell imaging was performed on a Zeiss LSM780 on a Zeiss Observer Z1 equipped with a Coherent Chameleon laser. Laser injury was performed using 100% of the Coherent Chameleon laser at 800 nm. Images were taken every 1.95 s. Video analysis was carried out using ImageJ software, and the Ca^2+^ peak intensity was determined by the quantification of fluorescence intensity in ten randomly chosen cells before and after laser damage per imaged field. The percentage of cells in the wave was assessed by counting the number of cells that had an increase in fluorescence intensity following laser injury in relation to the total number of cells per imaged field. The Fluo-3 transmission wave angle was assessed using the ImageJ ‘angle tool’. The number of cells that transmitted the Fluo-3 signal was determined in five independent experimental repeats.

### In vivo experiments

Animals were housed with free access to food and water in a 12h light/dark cycle. For survival experiments, Ragγ2C-/- mice were euthanized if they exhibited signs of neurological morbidity or if they lost > 20% of their body weight. Craniotomy and GBM spheroid implantation were done as previously described [24]. Briefly, a 5-mm circle was drilled between lambdoid, sagittal, and coronal sutures of the skullon ketamine/xylazine anesthetized mice. A 250-μm diameter P3 GBM cells spheroid was injected in the cortex and sealed with a glass coverslip cemented on top of the mouse skull. For multiphoton excitation of endogenous fluorophores in experimental gliomas, we used a Leica SP8 DIVE in vivo imaging system equipped with 4tune spectral external hybrid detectors and an InSightX3 laser (SpectraPhysics). The microscope was equipped with in house designed mouse holding platform for intravital imaging (stereotactic frame, Narishige; gas anesthesia and body temperature monitoring/control, Minerve). GFP signal from genetically modified tumor cells was acquired at 925-nm wavelength. Alexa Fluor 647 coupled Dextran was acquired at 1200-nm wavelength.

### Statistics

All statistical analyses were performed on GraphPad Prism 8.1.2. Data are displayed as mean ±SD. To compare one variable between multiple groups, one-way ANOVA with Tukey’s post hoc test was used. To compare 2 or more variables between multiple groups, two-way ANOVA with Tukey’s post hoc test was used. Differences were considered statistically significant when the p-value was below 0.05.

The difference in total MT length between responders/non-responders was analyzed using a T-test. Exploring the link between pSMAD3 expression and total MT length was done using a linear mixed model with compound symmetric correlation matrix. Model selection was done using the likelihood ratio test, first on the random effects using REML estimation, then on the fixed effects using ML estimation. The final model was estimated with REML. Statistical analysis was done using R software (Rstudio version 1.4.1103, R version 4.0.3). The linear mixed model was fitted using the nlme package (version 3.1). Semi-partial coefficient of determination R^2^ was calculated using the r2glmm package (version 0.1.2) [14].

## RESULTS

### TGF-β signaling is upregulated in GBM compared to 1p/19q co-deleted tumors

MTs are abundant in GBM, however uncommon in 1p/19q co-deleted oligodendroglioma. To identify new drivers of MT formation, we compared gene expression data from TCGA of IDH wild-type (wt) tumors, the majority being GBM, to IDH-mutant and 1p/19q co-deleted oligodendroglioma. We found that *TGFB1* and *TGFB2* were among the significantly differentially expressed genes (Supplementary Table 1). Both, *TGFB1* and *TGFB2* are highly upregulated in IDH-wt tumors compared to IDH-mutant and 1p/19q co-deleted tumors (Fig. 1a), and also upregulated in non-codeleted (IDH-wt and IDH-mutant astrocytoma/GBM) vs. co-deleted tumors (Supplementary Fig. 1a). MT formation has been previously observed in particular at the invasive front of GBM [26]. As TGF-β is a known driver of GBM invasion, we performed more detailed analyses. Interestingly, *TGFB1* is located on chromosome 19q, which makes it an attractive candidate for MT formation. The differentially expressed genes of IDH-wt compared to IDH-mutant and 1p/19q co-deleted tumors (1. comparison) were then used to conduct a Gene Ontology (GO) term enrichment analysis. The top regulated pathways were related to extracellular matrix, which are important pathways in tumor invasion (Fig. 1b). We performed the same GO term analysis for non-co-deleted versus co-deleted tumors (2. comparison) where extracellular matrix pathways were also found, however, below a number of immune-related pathways (Supplementary Fig. 1b). We then analyzed the commonly regulated pathways from these two comparisons in a Venn diagram revealing that the top common pathways in both were related to extracellular matrix (Supplementary Fig. 1 c,d). As MT formation is also highly dependent on extracellular matrix reorganization this could indicate an important role of TGF-β in this process. When focusing on the top 20 GO terms where either *TGFB1* or *TGFB2* was present in the gene list, the majority of associated genes in these pathways were upregulated in IDH-wt compared to 1p/19q co-deleted tumors (Fig. 1c). Next, we identified TGF-β responsive genes from the literature and showed that the majority of these genes are upregulated in IDH-wt compared to 1p/19q co-deleted tumors (Fig. 1d). Using connectivity map from the Broad institute [18], we analyzed drug candidates that would inhibit the expression signature of GBM (top 20 upregulated genes) compared to oligodendroglioma. One of the top drug candidates that came up in this screen were TGF-β inhibitors (Fig. 1e) further highlighting an important role of the TGF-β pathway in promoting a GBM gene expression signature, when compared to chemosensitive oligodendroglioma.

**Fig. 1.**
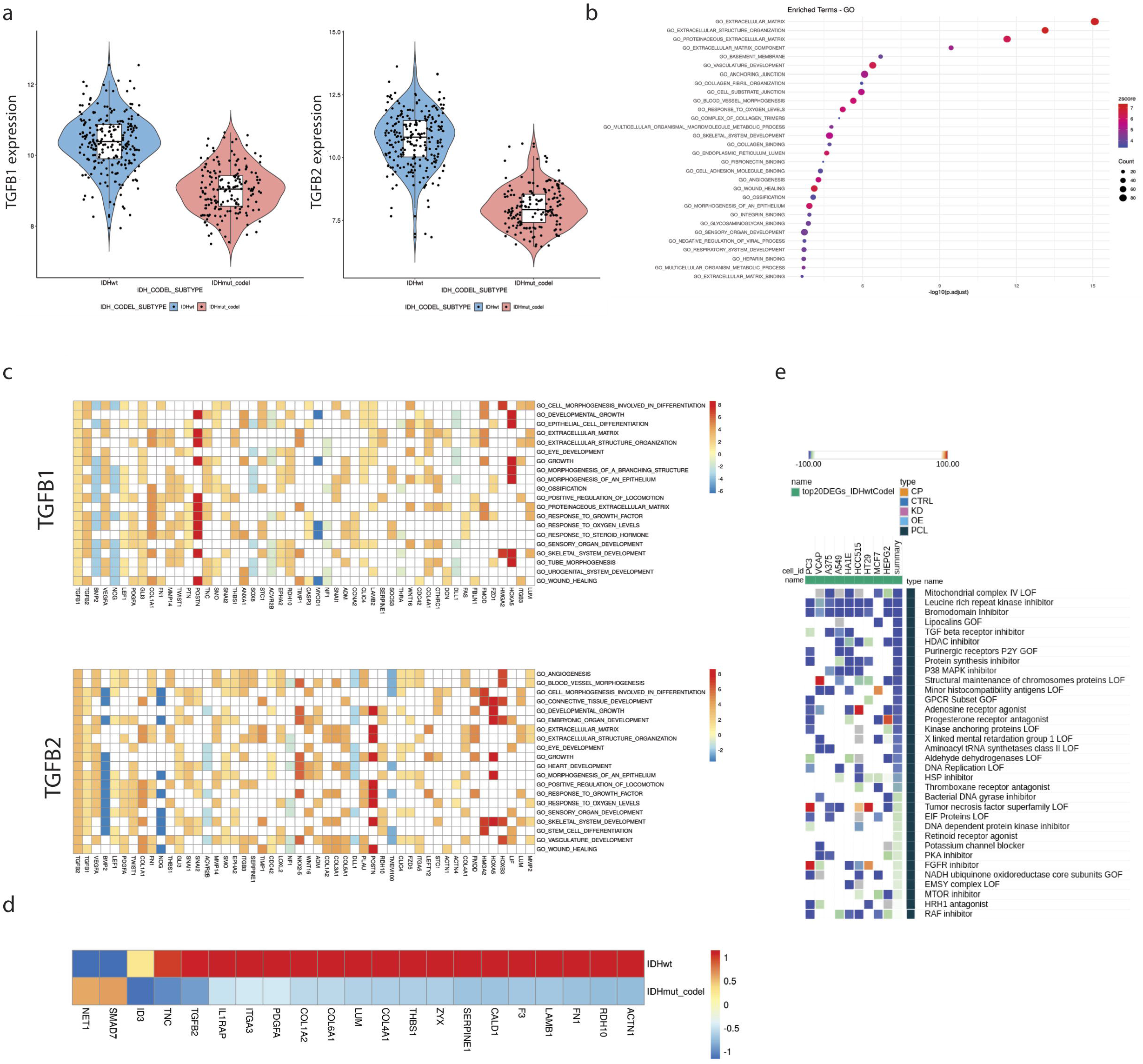
TGF-β signaling is upregulated in GBM compared to IDH-mutant and 1p/19q co-deleted oligodendroglioma. Analysis of TCGA data comparing IDH-wt GBM and IDH-mutant and 1p/19q co-deleted oligodendroglioma. **a** TGFB1 and TGFB2 are upregulated in IDH-wt tumors. **b** GO term analysis reveals pathways related to extracellular matrix. **c** Heatmap of genes related to the top 20 GO terms where either *TGFB1* or *TGFB2* is present. **d** TGF-β responsive genes identified from literature. **e** Analysis of drugs related to gene expression signature by connectivity map reveals that TGF-β inhibitors are one of the top candidate drugs that inhibit IDH-wt GBM signature.

### TGF-β promotes MT formation in GBM cell lines in vitro

To analyze the ability of different growth factors, including TGF-β, to induce cytoplasmic protrusions (MT-like structures), we stimulated P3 GBM cells with EGF, FGF, PDGFbb, TGF-β1 or TGF-β2. We observed that TGF-β1 was most efficient in promoting cellular protrusions when analyzing % of cells with protrusions, number of protrusions per cell and length of protrusions (Fig. 2a, Supplementary Figure 2a). Next, we stimulated 2 different GBM stem cell (GSC) lines (P3 and GG16) and 2 serum cultured GBM cell lines (U87 and U251) with TGF-β1 and analyzed morphological changes of the cells. A clear promotion of cellular protrusions by TGF-β1 was observed in all cell lines (Fig. 2b, Supplementary Fig. 2b,c). To verify that these protrusions are MTs we stained the cells for GAP43, a known driver for MT formation as well as nestin and actin to visualize the cytoskeleton. The protrusions were rich in GAP43 which colocalized with actin and nestin (Fig. 2b, Supplementary Fig. 2b). This indicated that the protrusions are MTs as described previously [26, 36]. To further verify that cells are interconnected through MTs upon TGF-β1 stimulation, we performed Electron microscopy. We observed MTs extending from one cell and inserting into the membrane of a neighboring cell (Fig. 2c). We analyzed and quantified the formation of MTs in detail by confocal microscopy using GAP43/nestin or actin stainings. TGF-β1 stimulation increased the number of cells connected through MTs and importantly, also the length of MTs increased significantly (Fig. 2d; Supplementary Fig. 2b,c). To block TGF-β signaling, we used the TGF-β inhibitor LY2157299, which significantly reduced pSMAD2 phosphorylation in P3 GBM cells (Supplementary Fig. 3a). MT formation was significantly inhibited by LY2157299: the number of MTs per cell, the number of cells with MTs and the length of MTs decreased substantially under treatment with LY2157299 in both P3 and GG16 GBM cells (Fig. 2d; Supplementary Fig. 2c and 3b). In addition, we used an inducible shRNA to knockdown TGFBRII, as stable knockdown was lethal to the cells (data not shown). The inducible construct confirmed the inhibition of MT formation in P3 cells (Supplementary Fig. 3c,d). As GAP43 has been shown to be a major structural element of MTs, we knocked down GAP43 with a shRNA to analyze if MT formation is blocked under TGF-β1 stimulation. As expected, GAP43 knockdown prevented a significant increase in the number and length of MTs under TGF-β1 stimulation (Supplementary Fig. 4) verifying that also in our culture system MT formation is dependent on GAP43 as described previously [26]. To demonstrate that there is a functional connectivity between the tumor cells through the MT network, we performed calcium imaging using fluorescence indicator of intracellular calcium 3 (Fluo-3) [25]. Stimulation with TGF-β1 resulted in increased Calcium exchange between tumor cells through the MT network which was inhibited by LY2157299 (Fig. 2e, Movies 1-4).

**Fig. 2.**
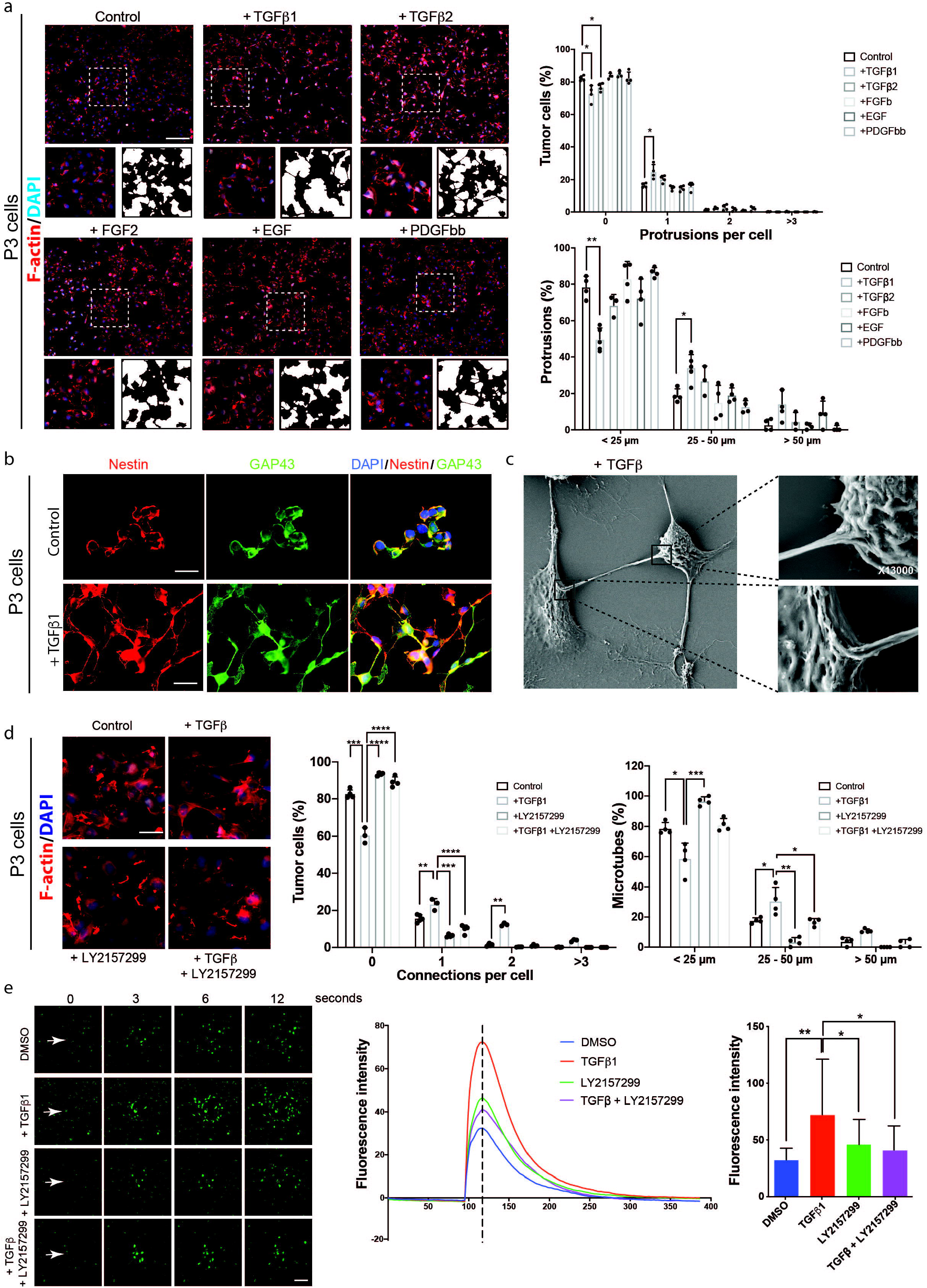
TGF-β promotes MT formation and communication via Calcium signaling in GBM cells. **a** TGF-β1 is the strongest inducer of cellular protrusions among a panel of different growth factors in P3 GBM cells. Immunofluorescence staining for F-actin is shown. Inserts are higher magnifications and black and white pictures to better visualize the MT network. Quantification of protrusion number and length is provided. Scale bar 50 µm. *p< 0.05; **p<0.01. **b** Cellular protrusions induced by TGF-β1 in P3 GBM cells are identified as MTs due to expression of GAP43 which co-localizes with the cytoskeleton protein nestin. Scale bar 20 µm. **c** Scanning Electron Microscopy of P3 GBM cells shows that MTs connect two neighboring cells through cytoplasmic insertions. Higher magnifications of specific areas are provided as indicated. **d** TGF-β inhibitor LY2157299 inhibits MT formation in P3 GBM cells. Immunofluorescence staining for F-actin is shown. Quantification of connections per cells and MT length is presented. Scale bar 10 µm. *p<0.05; **p<0.01; *** p<0.001. **e** Calcium exchange between tumour cells is significantly increased upon TGF-β stimulation and inhibited by LY2157299 in P3 GBM cells. Fluorescence intensity represents intensity of the Calcium signal. The images were taken seconds following the laser injury as indicated. The bar graph represents the intensity at the time point as indicated by the dotted line in the curve diagram. Scale bar 30 µm. *p<0.05; **p<0.01.

### TGF-β induced MT formation is associated with invasion

Among the GBM cell lines used in this work, we identified one GBM stem cell line, GG6, that did not show increase in MT formation upon TGF-β1 stimulation. GG16, used as a control cell line, confirmed significant MT formation under stimulation (Fig. 3a). As MT formation is associated with invasion, we aimed to analyze invasiveness of the non-responder cell line GG6 in comparison with the TGF-β1 responding GBM cell line GG16. We used a co-culture system of tumor spheroids with brain organoids from rat fetal brains to mimic the invasive process as described previously [6]. Invasion was quantified as indicated in methods. The non-responding cell line GG6 showed no significant invasion over a time period of 72h when co-cultured with brain organoids. In contrast, the TGF-β1 responding cell line GG16 showed substantial invasion into the organoid, which was not further increased by TGF-β1 stimulation, however significantly reduced by TGF-β inhibition with LY2157299, indicating endogenous presence of TGF-β in the organoid microenvironment (Fig. 3b). Upon implantation of both cell lines *in vivo*, we verified a less invasive phenotype of the non-responder GG6 compared to the responder GG16 and a reduced MT density in nestin immunostained sections (Fig. 3c). Importantly, we confirmed a reduced MT density in corresponding patient biopsies from GG6 compared to GG16 stained with nestin antibodies (Fig. 3c).

**Fig. 3.**
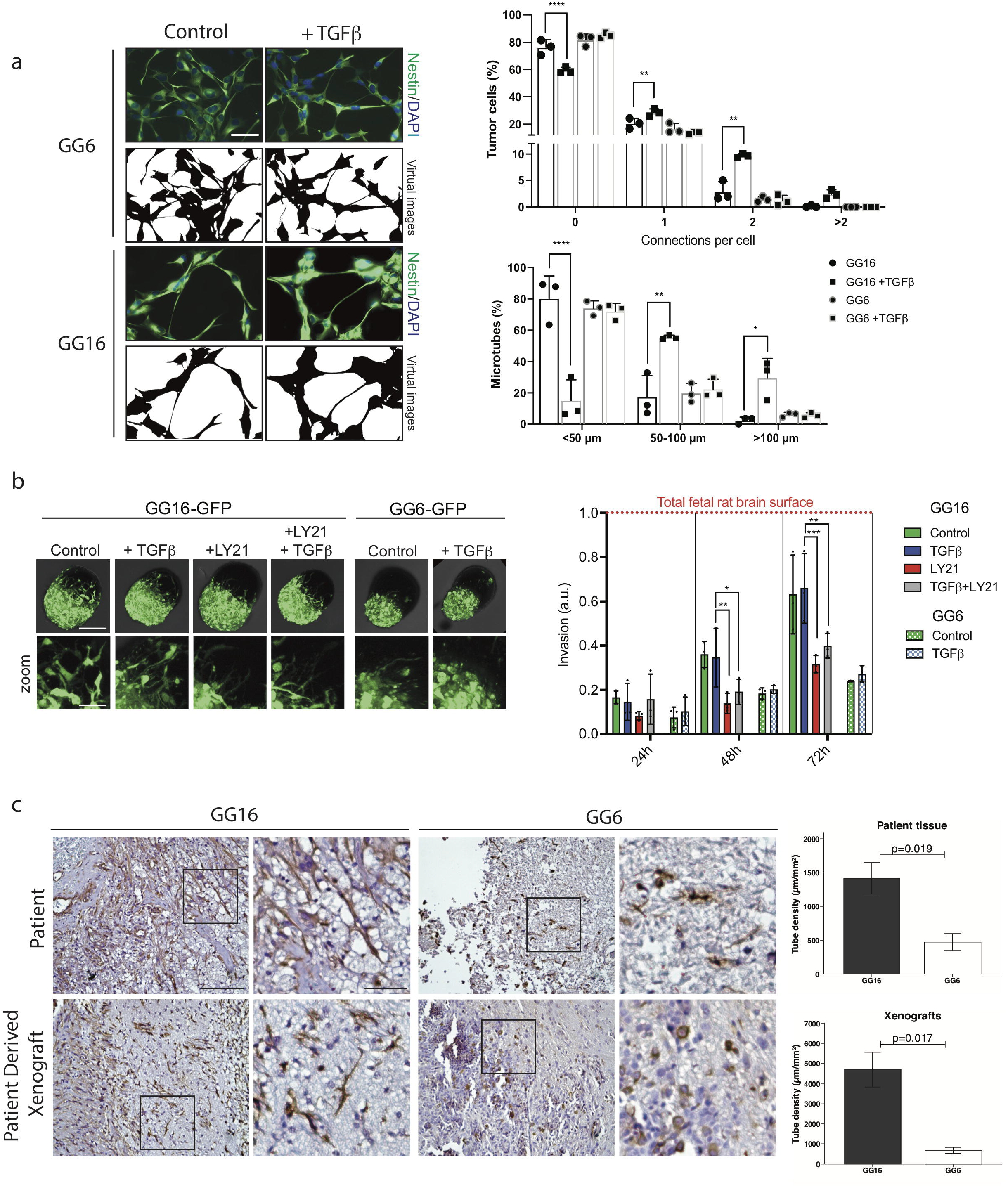
TGF-β induced MT formation is associated with invasion. **a** GG6 GBM cells are not responding to TGF-β1 stimulation with increased MT formation in contrast to GG16 GBM cells. Immunofluorescence stainings for nestin are shown. Black and white pictures are presented to better visualize the MT network. Quantification of connections per cell and MT length is presented. Scale bar 20 µm *p<0.05; **p<0.01; ***p0.001<; ****p<0.0001. **b** Invasion of GG16 GBM cells into brain organoids is inhibited by LY2157299. TGF-β stimulation does not significantly increase invasion most likely due to presence of TGF-β in the microenvironment. In contrast, non-responder GG6 GBM cells do not show significant invasion into brain organoids. Immunfluorescence pictures of organoids after 72 h showing GFP signal (GBM cells). Higher magnifications of the invasive areas are provided below each organoid picture. Quantification of invasive cells is provided after 24h, 48h and 72h. Scale bar 70 µm (zoom 14 µm). *p<0.05; **p<0.01; ***p<0.001. **c** Immunohistochemical staining of GG6 and GG16 patient and xenograft GBM with nestin antibodies showing an extensive MT network in GG16, which is low to absent in GG6. Quantification of MT density is provided. Scale bar 100 µm (zoom 10 µm).

### SMAD activation is important for MT formation

To identify major signaling mediators in TGF-β1 induced MT formation, we analyzed activation of downstream pathways under stimulation with TGF-β1 in P3 and GG16 cell lines. Only canonical (SMAD2/3) signaling was induced upon TGF-β1 stimulation, while non-canonical signaling (MAPK and Pi3K/Akt) was constitutively activated and did not increase upon TGF-β1 stimulation (Fig. 4a). Likewise, SMAD inhibition under TGF-β1 stimulation using the SMAD inhibitor SIS3, significantly reduced MT formation (Fig. 4b). To further substantiate the role of SMAD2/3 phosphorylation in MT formation, we immunostained consecutive sections from 7 GBM patient biopsies with pSMAD3 and nestin antibodies (Fig. 4c, Supplementary Fig. 5). Quantification of MT density in randomly selected areas was performed as described in the methods. For statistical analysis of the data, model selection was performed as indicated in the methods and supplementary table 2. The final model indicated a significant association between increased pSMAD3 expression and increased MT density when adjusting for overall cell density (T=3.29, p=0.0013).

**Fig. 4.**
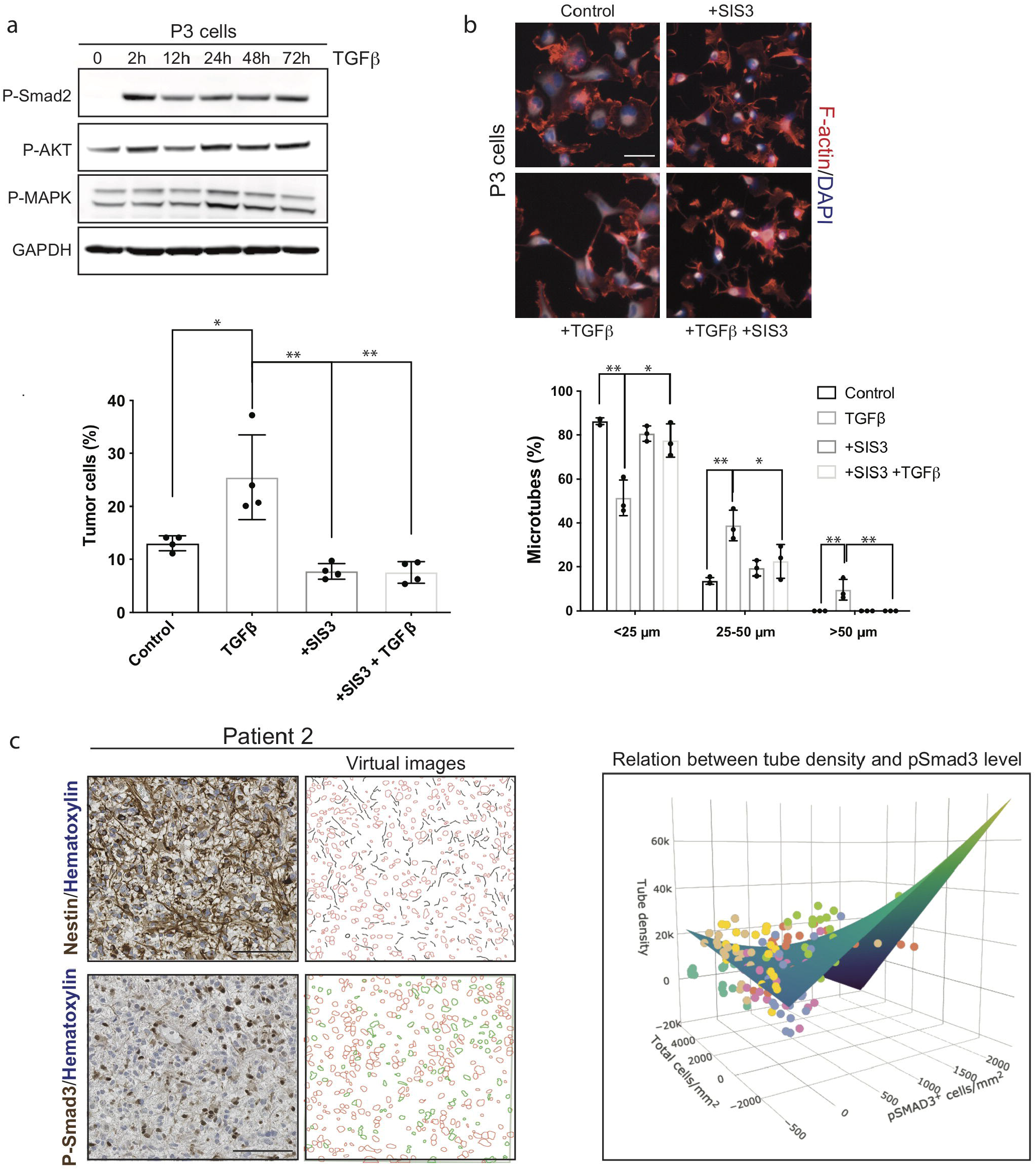
SMAD activation is important for MT formation. **a** Western blot for canonical TGF-β downstream signaling protein pSMAD2 and non-canonical proteins pMAPK and pAkt in P3 GBM cells with and without TGF-β1 stimulation. **b** SMAD inhibitor SIS3 inhibits MT formation under TGF-β1 stimulation. Immunofluorescence staining for F-actin is shown. Quantification of cells with MTs and MT length is presented. Scale bar 10 µm. *p<0.05; **p<0.01. **c** SMAD3 phosphorylation correlates with MT formation in GBM patient biopsies. Quantification of MT length on consecutive pSMAD3 immunostained and nestin immunostained sections is shown (see methods for details). The graph shows correlation of pSMAD3 expression with MT length. The results of all patients are included in the graph. Scale bar 100 µm; p=0.0013.

### TSP1 is a candidate for MT formation downstream of TGF-β**1 and Smad2/3**

To identify targetable proteins involved in MT formation downstream of TGF-β1 and SMAD activation, we performed RNA sequencing of the responder cell lines GG16 and P3 and the non-responder cell line GG6 with and without TGF-β1 stimulation (48h) to identify differences in gene expression profiles that would indicate potential candidates for MT formation downstream of TGF-β1 /SMAD2/3 activation. The number of significantly upregulated genes under TGF-β1 stimulation varied substantially between the cell lines (Supplementary Table 3). In particular GG16 showed only 60 upregulated genes, however, 34 of these genes were shared with P3, while only 1 gene was shared with the non-responder cell line GG6 (Fig. 5a, Supplementary Table 3). 19 genes were commonly upregulated in all 3 cell lines. Gene enrichment analysis showed that in all 3 cell lines, extracellular structure organization and extracellular matrix organization were the top regulated GOs (Fig. 5b). When analyzing these GOs for common genes between the responders P3 and GG16, only 4 genes were found, while also 4 genes were in common between all three cell lines (Fig. 5c, Supplementary tables 4 and 5). Among the 4 genes in common between the responders P3 and GG16, *TSP1* (*THBS1*) was identified as an interesting candidate because it also showed up in the previous analysis of TCGA data (Fig. 1 c,d). Moreover, we have recently reported that TSP1 is a downstream mediator of TGF-β1 and SMAD3 signaling in GBM and important in tumor cell invasion [10]. Analysis of TCGA data also confirmed upregulation of TSP1 in IDH-wt GBM compared to IDH-mutant and 1p/19q co-deleted oligodendroglioma (Fig. 5d).

**Fig. 5.**
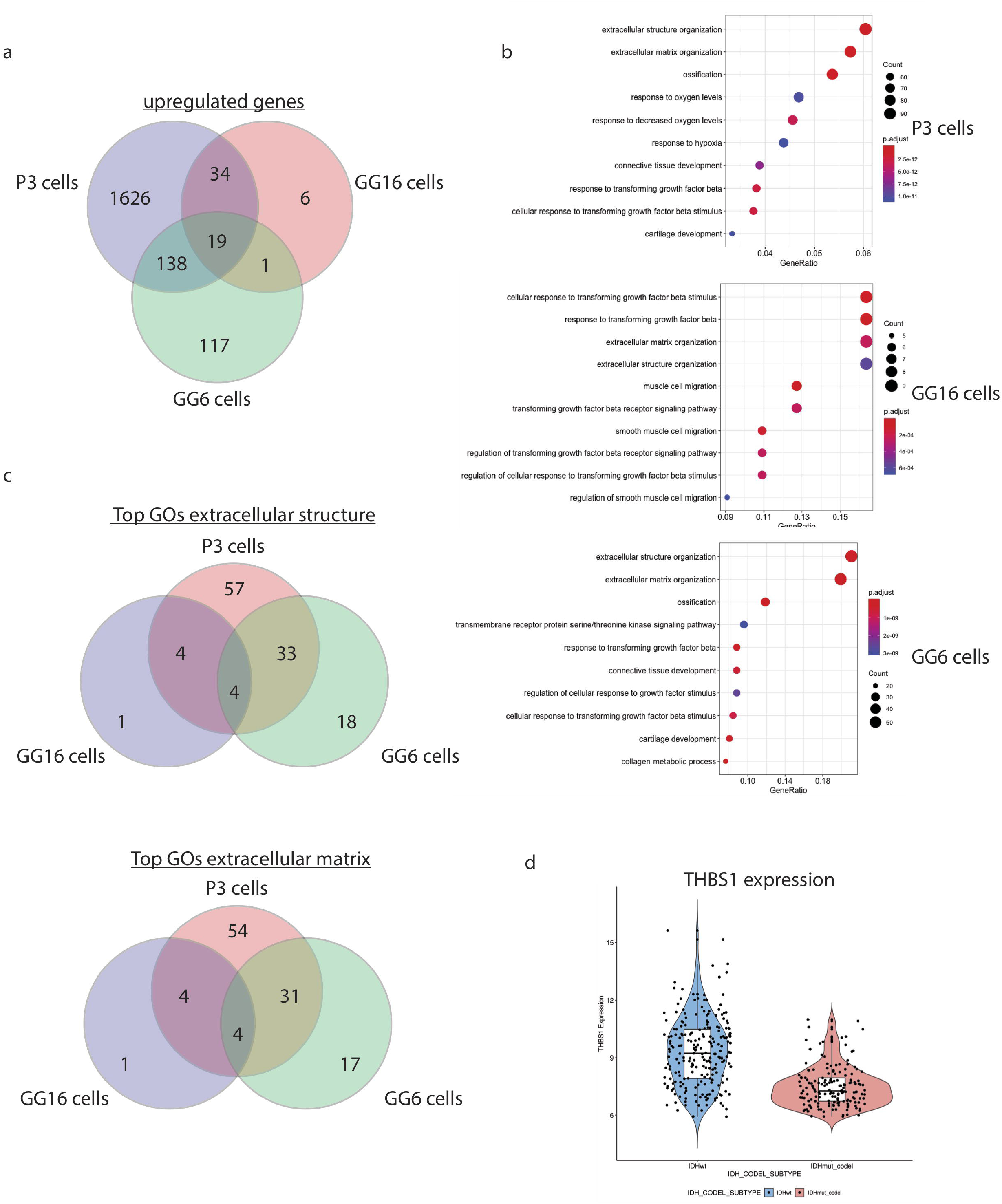
TSP1 is a candidate for MT formation downstream of TGF-β1 and Smad2/3. RNA sequencing data of P3, GG6 and GG16 GBM cells unstimulated and stimulated with TGF-β1 for 48h. **a** Venn diagram showing number of common and unique upregulated genes upon TGF-β1 stimulation. **b** Gene enrichment analysis showing upregulation of pathways related to extracellular matrix/structure in all 3 cell lines. **c** Venn diagram showing number of common and unique regulated genes among the 3 cell lines from extracellular matrix and extracellular structure GOs. **d** Comparison of TSP1 expression in TCGA data between IDH-wt and IDH-mutant, 1p/19q co-deleted tumors.

### TSP1 is activated by TGF-β1 and mediates MT formation

As described previously [10], Tsp1 expression is upregulated by TGF-β signaling through activation of the Tsp1 promoter by pSMAD3 (Figure 6a). Here, we confirmed that TSP1 protein expression was induced upon TGF-β1 stimulation in the responder cell lines GG16 and P3, while it was absent in the non-responder cell line GG6 (Fig. 6b; Supplementary Figure 6). We used a shRNA to knock down expression of TSP1 and analyzed MT formation under TGF-β1 stimulation. ShTsp1 significantly reduced MT formation (connections per cell and MT length) when compared to a scrambled control (Fig. 6c).

**Fig. 6.**
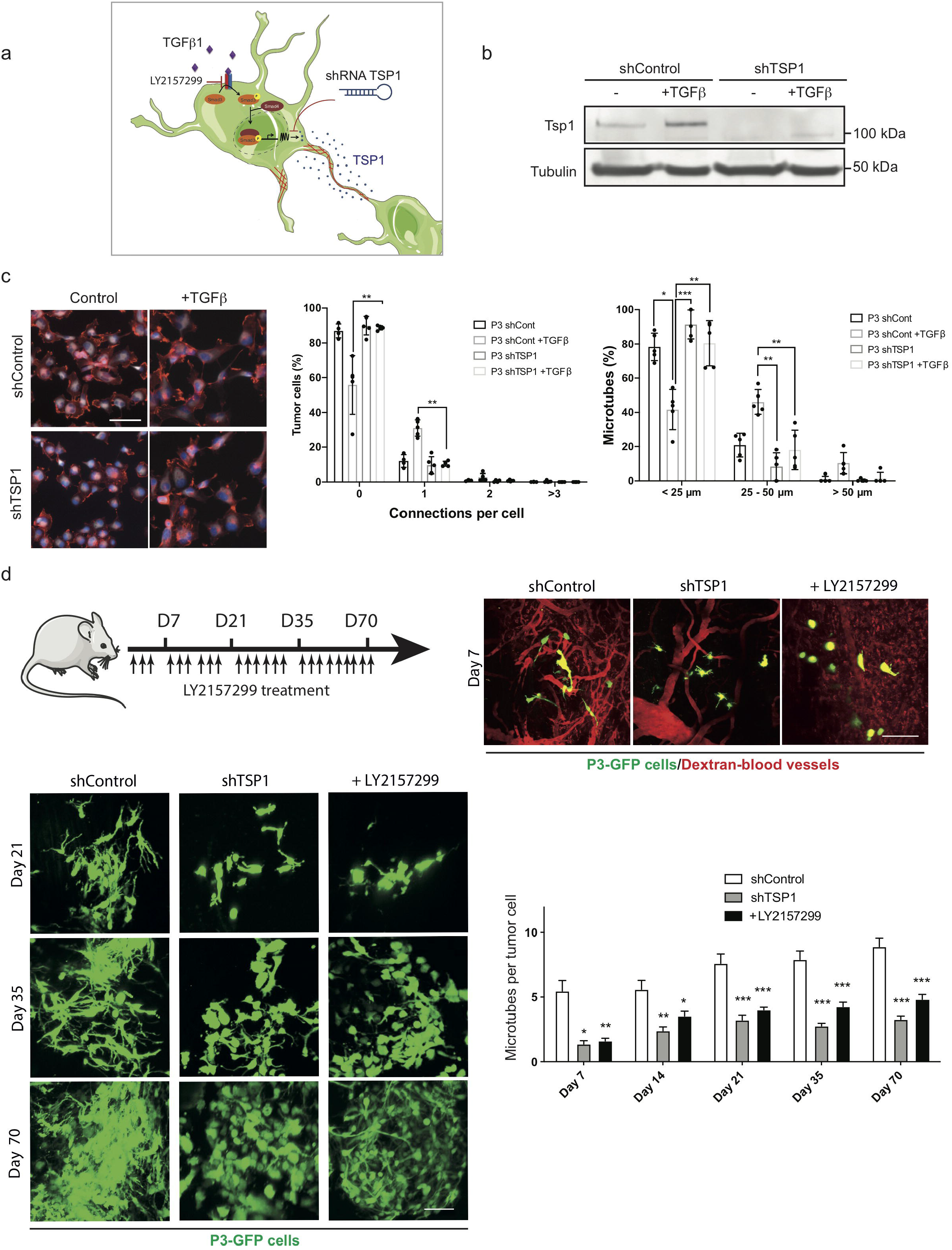
Inhibition of TSP1 reduces MT formation *in vitro* and *in vivo*. **a** Schematic figure showing upregulation of Tsp1 expression by TGF-β signaling through binding of pSMAD3 to the Tsp1 promoter. Tsp1 expression can be inhibited by LY2157299 or shRNA Tsp1. **b** Western blot of Tsp1 in P3 shControl and P3 shTsp1 with and without TGF-β1 stimulation. **c** MT formation is inhibited by shTsp1. Immunfluorescence staining for F-actin is shown. Quantification of connections per cell and MT length is presented. Scale bar 10 µm. *p<0/05; **p<0.01; ***p<0.001. **d** Formation of MT network in vivo in the orthotopic P3-GFP GBM xenograft model was analyzed by intravital imaging. ShTsp1 and LY2157299 inhibit MT formation compared to shcontrol. Immunofluorescence pictures of intravital imaging are shown. Quantification of MT number per tumour cell is presented. Scale bar 150 µm. *p<0.05; **p<0.01; ***p<0.001.

To verify that TGF-β and TSP1 are important players of MT formation in vivo, we performed intravital imaging of P3 cells implanted orthotopically in NOD/SCID mice. After 7 to 70 days both, inhibition of TGF-β by LY2157299 and knockdown of TSP1 by shRNA reduced the number of MTs per tumour cell *in vivo* compared to controls (Fig. 6d).

## Discussion

In the present study we identified a new molecular mechanism of MT formation in GBM. We showed that TGF-β1, a multi-functional cytokine, promotes MT formation *in vitro* and *in vivo*. MTs have been defined as major drivers of GBM invasion and resistance to standard treatments such as radiation and chemotherapy [26]. Since this hallmark study, where the GAP43 protein was identified as a major structural protein of MTs, new and fundamental insights into detailed molecular mechanisms of MT formation are still lacking. In search of molecular drivers of MT formation, we performed a bioinformatics analysis comparing IDH-mutant 1p/19q co-deleted versus IDH-wt tumors. We chose this starting point because MT formation is abundant in IDH-wt tumors while it is low to absent in IDH-mutant and co-deleted tumors [26]. This analysis brought our attention to TGF-β. Among other gene candidates, TGFB1 and TGFB2 are expressed at significantly lower levels in co-deleted tumors. Importantly, *TGFB1* is located on chromosome 19q and very important in GBM invasion [8], which are additional justifications for its potential role in MT formation. TGF-β is a cytokine with multiple functions which are dependent on the cell type and its associated microenvironment [4]. In GBM, TGF-β is well characterized as a cytokine that promotes invasion, angiogenesis as well as suppression of the immune system [5]. Thus, our results add an additional function of TGF-β to its divergent roles in GBM development. We demonstrated experimentally in different ways that TGF-β1 is important in MT formation: 1. by stimulation with TGF-β1; 2. by blocking of TGF-β activity using the inhibitor LY2157299 and an inducible shRNA. As our study is among the first to analyze MT formation in vitro, we had to verify that the structures we analyzed are corresponding to the MTs which have been shown in vivo [26]. We identified MTs in our culture system by demonstrating co-expression of major structural proteins of MTs such as nestin, actin and GAP43. Nestin, a protein related to cellular stemness, has recently been identified as an important protein expressed in the GBM MT network *in vivo* [36]. Thus, we used this marker in our study to identify and quantify MTs. Knockdown of GAP43, which significantly reduced MT formation under TGF-β1 stimulation, further corroborated that the structures we analyzed *in vitro* are very similar to the MTs formed *in vivo*.

MTs connect tumor cells with each other forming a communicating network. To show that network formation is promoted by TGF-β1, we performed Calcium imaging. As expected, Calcium exchange increased upon TGF-β1 stimulation and was inhibited by LY2157299. Network formation and increase in Calcium exchange in GBM cells *in vitro* was also observed by da Silva et al. using ROCK inhibition [9].

The majority of our cell lines responded to TGF-β1 with increase in MT formation. Interestingly, we identified one cell line (GG6), which did not respond and also showed very limited invasive capacity upon TGF-β stimulation using *ex vivo* brain organoid co-cultures and *in vivo* xenografts. These results highlight that MT formation and invasion are highly connected as also demonstrated previously [26]. Inhibition of TGF-β activity in GBM in a clinical setting has been unsuccessful so far [7, 35]. Reasons for these disappointing results may be the dose limitations due to the important physiological roles of TGF-β in the body and also the BBB which often impairs drug penetration [27]. Thus, we aimed to further characterize downstream signaling of TGF-β inducing MT formation to reveal more cancer-specific downstream signaling. First, we identified the canonical SMAD2/3 signaling in being the most important downstream pathway and also showed in a patient setting that SMAD3 activation correlates with MT length in patient biopsies. Previously, SMAD2/3 signaling was shown to impact glioma invasion[16, 21], whereas SMAD3 was defined as the main transcription factor regulating invasion and TSP1 expression in GBM cells [10]. To further identify more specific downstream targets of TGF-β and SMAD signaling, we performed a Bioinformatics analysis of cell lines P3 and GG16 (responding to TGF-β with MT formation) and compared the data to the non-responder cell line GG6. TSP1 came up as an interesting candidate which was absent in the non-responder GG6 as also verified by western blotting. Interestingly, we have previously shown that TSP1 is activated by TGF-β signaling and specifically by SMAD3 through a SMAD3 binding site in the TSP1 promoter [10]. We also demonstrated in this study that TSP1 promotes GBM invasion downstream of TGF-β and SMAD. Thus, as invasion and MT formation are linked to each other, we demonstrated that knockdown of TSP1 inhibited invasion and MT formation both *in vitro* and *in vivo*.

In conclusion, we identified TGF-β as a new mediator of MT formation through SMAD and Tsp1 signaling. Blocking this pathway might be an attractive strategy to treat GBM in the future.

## Acknowledgments

We thank B. Nordanger and H. S. Sdik for expert technical assistance and the Molecular Imaging Center (MIC) in Bergen, Norway for technical support. Simon Storevik was supported by a fellowship from Helse Vest. This work was supported by the Norwegian Cancer Society, Fondation ARC, Ligue Contre le Cancer and ARTC.

## Notes

### Competing Interest Statement

The authors have declared no competing interest.

### Summary of Updates

author list updated

## References

1 Adams JC, Lawler J (2011) The thrombospondins. Cold Spring Harb Perspect Biol 3: a009712 Doi 10.1101/cshperspect.a009712

2 Anders S, Huber W (2010) Differential expression analysis for sequence count data. Genome Biol 11: R106 Doi 10.1186/gb-2010-11-10-r106

3 Arganda-Carreras I, Fernandez-Gonzalez R, Munoz-Barrutia A, Ortiz-De-Solorzano C (2010) 3D reconstruction of histological sections: Application to mammary gland tissue. Microsc Res Tech 73: 1019–1029 Doi 10.1002/jemt.20829

4 Batlle E, Massague J (2019) Transforming Growth Factor-beta Signaling in Immunity and Cancer. Immunity 50: 924–940 Doi 10.1016/j.immuni.2019.03.024

5 Birch JL, Coull BJ, Spender LC, Watt C, Willison A, Syed N, Chalmers AJ, Hossain-Ibrahim MK, Inman GJ (2020) Multifaceted transforming growth factor-beta (TGFbeta) signalling in glioblastoma. Cell Signal 72: 109638 Doi 10.1016/j.cellsig.2020.109638

6 Bjerkvig R, Laerum OD, Mella O (1986) Glioma cell interactions with fetal rat brain aggregates in vitro and with brain tissue in vivo. Cancer Res 46: 4071–4079

7 Brandes AA, Carpentier AF, Kesari S, Sepulveda-Sanchez JM, Wheeler HR, Chinot O, Cher L, Steinbach JP, Capper D, Specenier P et al (2016) A Phase II randomized study of galunisertib monotherapy or galunisertib plus lomustine compared with lomustine monotherapy in patients with recurrent glioblastoma. Neuro Oncol 18: 1146–1156 Doi 10.1093/neuonc/now009

8 Caja L, Bellomo C, Moustakas A (2015) Transforming growth factor beta and bone morphogenetic protein actions in brain tumors. FEBS Lett 589: 1588–1597 Doi 10.1016/j.febslet.2015.04.058

9 da Silva B, Irving BK, Polson ES, Droop A, Griffiths HBS, Mathew RK, Stead LF, Marrison J, Williams C, Williams J et al (2019) Chemically induced neurite-like outgrowth reveals a multicellular network function in patient-derived glioblastoma cells. J Cell Sci 132: Doi 10.1242/jcs.228452

10 Daubon T, Leon C, Clarke K, Andrique L, Salabert L, Darbo E, Pineau R, Guerit S, Maitre M, Dedieu S et al (2019) Deciphering the complex role of thrombospondin-1 in glioblastoma development. Nat Commun 10: 1146 Doi 10.1038/s41467-019-08480-y

11 Durinck S, Spellman PT, Birney E, Huber W (2009) Mapping identifiers for the integration of genomic datasets with the R/Bioconductor package biomaRt. Nat Protoc 4: 1184–1191 Doi 10.1038/nprot.2009.97

12 Eskilsson E, Rosland GV, Talasila KM, Knappskog S, Keunen O, Sottoriva A, Foerster S, Solecki G, Taxt T, Jirik R et al (2016) EGFRvIII mutations can emerge as late and heterogenous events in glioblastoma development and promote angiogenesis through Src activation. Neuro Oncol 18: 1644–1655 Doi 10.1093/neuonc/now113

13 Firlej V, Mathieu JR, Gilbert C, Lemonnier L, Nakhle J, Gallou-Kabani C, Guarmit B, Morin A, Prevarskaya N, Delongchamps NB et al (2011) Thrombospondin-1 triggers cell migration and development of advanced prostate tumors. Cancer Res 71: 7649–7658 Doi 10.1158/0008-5472.CAN-11-0833

14 Jaeger B, Edwards L, Das K, Sen P (2016) An R2 statistic for fixed effects in the generalized linear mixed model. Journal of Applied Statistics 44: 1086–1105

15 Jerman T, Pernus F, Likar B, Spiclin Z (2015) Beyond Frangi: an improved multiscale vesselness filter. SPIE Medical Imaging. Proceedings Volume 9413, Medical Imaging 2015: Image Processing, City

16 Joseph JV, Conroy S, Tomar T, Eggens-Meijer E, Bhat K, Copray S, Walenkamp AM, Boddeke E, Balasubramanyian V, Wagemakers M et al (2014) TGF-beta is an inducer of ZEB1-dependent mesenchymal transdifferentiation in glioblastoma that is associated with tumor invasion. Cell Death Dis 5: e1443 Doi 10.1038/cddis.2014.395

17 Jung E, Osswald M, Blaes J, Wiestler B, Sahm F, Schmenger T, Solecki G, Deumelandt K, Kurz FT, Xie Ret al (2017) Tweety-Homolog 1 Drives Brain Colonization of Gliomas. J Neurosci 37: 6837–6850 Doi 10.1523/JNEUROSCI.3532-16.2017

18 Lamb J, Crawford ED, Peck D, Modell JW, Blat IC, Wrobel MJ, Lerner J, Brunet JP, Subramanian A, Ross KN et al (2006) The Connectivity Map: using gene-expression signatures to connect small molecules, genes, and disease. Science 313: 1929–1935 Doi 10.1126/science.1132939

19 Liao Y, Smyth GK, Shi W (2014) featureCounts: an efficient general purpose program for assigning sequence reads to genomic features. Bioinformatics 30: 923–930 Doi 10.1093/bioinformatics/btt656

20 Liberzon A, Subramanian A, Pinchback R, Thorvaldsdottir H, Tamayo P, Mesirov JP (2011) Molecular signatures database (MSigDB) 3.0. Bioinformatics 27: 1739–1740 Doi 10.1093/bioinformatics/btr260

21 Liu Z, Kuang W, Zhou Q, Zhang Y (2018) TGF-beta1 secreted by M2 phenotype macrophages enhances the stemness and migration of glioma cells via the SMAD2/3 signalling pathway. Int J Mol Med 42: 3395–3403 Doi 10.3892/ijmm.2018.3923

22 Louis DN, Perry A, Reifenberger G, von Deimling A, Figarella-Branger D, Cavenee WK, Ohgaki H, Wiestler OD, Kleihues P, Ellison DW (2016) The 2016 World Health Organization Classification of Tumors of the Central Nervous System: a summary. Acta Neuropathologica 131: 803–820 Doi 10.1007/s00401-016-1545-1

23 Love MI, Huber W, Anders S (2014) Moderated estimation of fold change and dispersion for RNA-seq data with DESeq2. Genome Biol 15: 550 Doi 10.1186/s13059-014-0550-8

24 Mathivet T, Bouleti C, Van Woensel M, Stanchi F, Verschuere T, Phng LK, Dejaegher J, Balcer M, Matsumoto K, Georgieva PB et al (2017) Dynamic stroma reorganization drives blood vessel dysmorphia during glioma growth. EMBO Mol Med 9: 1629–1645 Doi 10.15252/emmm.201607445

25 Minta A, Kao JP, Tsien RY (1989) Fluorescent indicators for cytosolic calcium based on rhodamine and fluorescein chromophores. J Biol Chem 264: 8171–8178

26 Osswald M, Jung E, Sahm F, Solecki G, Venkataramani V, Blaes J, Weil S, Horstmann H, Wiestler B, Syed M et al (2015) Brain tumour cells interconnect to a functional and resistant network. Nature 528: 93–98 Doi 10.1038/nature16071

27 Pandit R, Chen L, Gotz J (2019) The blood-brain barrier: Physiology and strategies for drug delivery. Adv Drug Deliv Rev: Doi 10.1016/j.addr.2019.11.009

28 Stupp R, Mason WP, van den Bent MJ, Weller M, Fisher B, Taphoorn MJ, Belanger K, Brandes AA, Marosi C, Bogdahn U et al (2005) Radiotherapy plus concomitant and adjuvant temozolomide for glioblastoma. N Engl J Med 352: 987–996 Doi 10.1056/NEJMoa043330

29 Team R (2016) RStudio: Integrated Development for R. RStudio, Inc, Boston, MA URL http://wwwrstudiocom/:

30 Team RC (2018) R: A language and environment for statistical computing.. R Foundation for Statistical Computing, Vienna https://wwwR-projectorg:

31 van den Bent MJ (2001) New perspectives for the diagnosis and treatment of oligodendroglioma. Expert Rev Anticancer Ther 1: 348–356 Doi 10.1586/14737140.1.3.348

32 Venkataramani V, Tanev DI, Strahle C, Studier-Fischer A, Fankhauser L, Kessler T, Korber C, Kardorff M, Ratliff M, Xie Ret al (2019) Glutamatergic synaptic input to glioma cells drives brain tumour progression. Nature 573: 532–538 Doi 10.1038/s41586-019-1564-x

33 Walter W, Sanchez-Cabo F, Ricote M (2015) GOplot: an R package for visually combining expression data with functional analysis. Bioinformatics 31: 2912–2914 Doi 10.1093/bioinformatics/btv300

34 Weil S, Osswald M, Solecki G, Grosch J, Jung E, Lemke D, Ratliff M, Hanggi D, Wick W, Winkler F (2017) Tumor microtubes convey resistance to surgical lesions and chemotherapy in gliomas. Neuro Oncol 19: 1316–1326 Doi 10.1093/neuonc/nox070

35 Wick A, Desjardins A, Suarez C, Forsyth P, Gueorguieva I, Burkholder T, Cleverly AL, Estrem ST, Wang S, Lahn MM et al (2020) Phase 1b/2a study of galunisertib, a small molecule inhibitor of transforming growth factor-beta receptor I, in combination with standard temozolomide-based radiochemotherapy in patients with newly diagnosed malignant glioma. Invest New Drugs: Doi 10.1007/s10637-020-00910-9

36 Xie R, Kessler T, Grosch J, Hai L, Venkataramani V, Huang L, Hoffmann DC, Solecki G, Ratliff M, Schlesner M et al (2020) Tumor cell network integration in glioma represents a stemness feature. Neuro Oncol: Doi 10.1093/neuonc/noaa275

37 Yu G, Wang LG, Han Y, He QY (2012) clusterProfiler: an R package for comparing biological themes among gene clusters. OMICS 16: 284–287 Doi 10.1089/omi.2011.0118

