## Supplementary Legends for "TGF-β promotes microtube formation in glioblastoma through Thrombospondin 1"

**Supplementary Figure Legends**

**Suppl. Figure 1: TGF-β signaling is upregulated in non-codeleted tumors compared to 1p/19q co-deleted oligodendroglioma.** **a** TGFB1 and TGFB2 are upregulated in non-codeleted tumors. **b** GO term analysis reveals pathways related to extracellular structure/matrix below a number of immune system related pathways. **c** Venn diagram showing number of unique and common pathways between the indicated comparisons. **d** common pathways from c are related to extracellular matrix/structure.

**Suppl. Figure 2:** **TGF-β promotes MT formation in GBM cells. a** TGF-β1 is the strongest inducer of cellular protrusions among a panel of different growth factors in P3 GBM cells. Quantification of tumour cells with protrusions is shown. *p<0.05; ***p< 0.001. **b** Cellular protrusions induced by TGF-β1 in the two serum-cultured cell lines U87 and U251 are identified as microtubes which express GAP43, nestin and actin. Quantification of connections per cell and MT length is presented. Scale bar 20 µm. *p<0.05; **p<0.01; *** p<0.001; **** p<0.0001. **c** TGF-β inhibitor LY2157299 inhibits microtube formation in GG16 GBM cells. Immunofluorescence staining for F-actin is shown. Quantification of connections per cell and MT length is presented. Scale bar 20 µm. *p<0.05; **p<0.01; **** p<0.0001.

**Suppl. Figure 3: TGF-β induced MT network is inhibited by LY2157299 and an inducible shRNA for TGFBRII. a** LY2157299 inhibits SMAD2 phosphorylation. Immunofluorescence staining with anti-actin and anti-pSMAD2 antibodies is shown. Scale bar 10 µm. **b** TGF-β inhibitor LY2157299 inhibits MT formation in P3 GBM cells. Quantification of tumour cells with MTs is presented. **c** western blot showing knockdown of TGFBRII with the inducible shTGFBRII. **d** shTGFBRII inhibits MT formation in P3 cells. Immunfluorescence staining for F-actin; Scale bar 10 µm. Quantification of tumour cells with MTs as well as MT length is shown. *p<0.05; **p<0.01; *** p<0.001.

**Suppl. Figure 4: shGAP43 inhibits the TGF-β induced MT network. a** Western blot of P3 transduced with shControl and P3 transduced with shGAP43 for GAP43 and Vinculin. **b** MT formation is inhibited by shGAP43 in P3 cells. Immunofluorescence staining with anti-actin and antibodies is shown; Scale bar 10 µm. Quantification of MT number and length is presented. *p<0.05; **p<0.01; *** p<0.001.

**Suppl. Figure 5: SMAD3 activation correlates with microtube length in patient biopsies. a** Two corresponding immunostained sections for nestin and pSMAD3 of GBM patient 2 are shown. Analyzed areas are marked as squares. **b** Correlation graphs for all patients used in this study.

**Suppl. Figure 6: Expression of Tsp1 is induced by TGF-β in GG16, but not in GG6.** Western blot of GG6 (Non-responder) and GG16 (Responder) for Tsp1 and Tubulin.

**Movie 1: Calcium exchange in P3 GBM cells (related to Figure 2e).** DMSO control.

**Movie 2: Calcium exchange in P3 GBM cells (related to Figure 2e).** TGF- β1.

**Movie 3: Calcium exchange in P3 GBM cells (related to Figure 2e).** LY2157299.

**Movie 4: Calcium exchange in P3 GBM cells (related to Figure 2e).** TGF- β1 + LY2157299.

**Legend for Supplementary table 2:**

Fixed effects are the average effect of predictor variables across patients. Their estimated regression coefficients - i.e. the average effect of a unit increase in a predictor on the outcome - is reported with confidence interval. The intercept represents the baseline tube density in an area with average pSMAD3 expression and cell density. R², the proportion of variance in tube density that can be explained by a predictor variable, is also reported with confidence interval for the fixed effects. The final model was a linear mixed model with fixed effects for pSMAD3+ cell density, total cell density and an interaction term; patient as the grouping variable; and finally random intercept and random slope for pSMAD3. The standard deviance reported for the random effects is a measure of the variation in intercept and slope between patients.
